# Biophotons: low signal/noise ratio reveals crucial events

**DOI:** 10.1101/558353

**Authors:** Maurizio Benfatto, Elisabetta Pace, Catalina Curceanu, Alessandro Scordo, Alberto Clozza, Ivan Davoli, Massimiliano Lucci, Roberto Francini, Fabio De Matteis, Maurizio Grandi, Rohisha Tuladhar, Paolo Grigolini

## Abstract

We study the emission of photons from germinating seeds using an experimental technique designed to detect photons of extremely small intensity when the signal/noise ratio is low. We analyze the dark count signal in the absence of germinating seeds as well as the photon emission during the germination process. The technique of analysis adopted here was originally designed to measure the temporal complexity of astrophysical, sociological and physiological processes. The foundation of this method, called Diffusion Entropy Analysis (DEA), rests on Kolmogorov complexity. The updated version of DEA used in this paper is designed to determine if the signal complexity is generated by either non-ergodic crucial events with a non-stationary correlation function or by the infinite memory of a stationary but non-integrable correlation function or by a mixture of both processes. We find that dark count yields the ordinary scaling, thereby showing that no complexity of either kinds may occur in the absence of any seeds in the chamber. In the presence of seeds in the chamber anomalous scaling emerges, reminiscent of that found in neuro-physiological processes. However, this is a mixture of both processes and with the progress of germination the non-ergodic component tends to vanish and complexity is dominated by the stationary infinite memory. We argue that this may be a sign of quantum coherence that according to some authors is the important ingredient of cognition.

## I. INTRODUCTION

An impressive revolution is occurring in biology [1, 2]. The traditional approach to Darwinian evolution based on the transmission of information through genes is integrated by epigenetic processes which rests instead on cell to cell communication and exchange of information between complex networks of organisms where each network of organisms represent a sort of social intelligence. In addition, the single cell itself is an intelligent system and the brain should not be interpreted as a supercomputer but rather as “an entire community of supercomputers ” [3]. Gordana Dodig-Crnkovic [4] points out that although until recently intelligence was considered a property of human beings, but now due to the autopoiesis of Maturana and Varela [5] cognition should be extended to all biological systems. This observation has the impressive effect of marking the breakdown of separation between different disciplines. There is, in fact, a growing evidence in psychology about the existence of a form of collective intelligence established by the dialogue between different people [6]. Thus, it seems that the discussion about cognition moves from the single cell molecular biology to a set of individuals, sociology and psychology, going through an intermediate bridge involving the brain of the single individual-neurophysiology. This generates two connected problems: (a) What is the origin of intelligence? (b) How do intelligent systems communicate?

As far as the origin of cognition is concerned, we find that in the literature, the predominant conjecture is that of Self Organized Criticality (SOC) [1, 2]. According to the illuminating arguments of Plenz [7], criticality plays a central role in the dynamics of the brain which was studied by the authors of [8]. The authors of [9] also show that criticality is not limited to the human brain but can be used to explain the behavior of a swarm of birds. The intelligence of a swarm of birds affords an intuitive way to acquaint the readers with the cognition that emerges from the germination of seeds as represented by the *crucial events*. In fact, the authors of Ref. [10] studied the phase-transition model of Vicsek *et al.* [11] and found that from time to time the flying speed of the swarm falls to virtually vanishing values making it possible for the swarm to select new flying directions. The time distance *τ* between two consecutive changes of flying directions is given by the waiting time distribution density *ψ*(*τ*) with the inverse power law structure

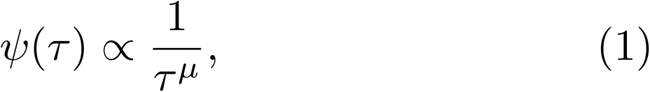

with *µ* = 1.35.

These abrupt changes of directions are a form of intermittence generated by criticality. Contoyiannis *et al*. [12] studied the criticality-induced intermittence of the Ising model which is in line with the theoretical arguments of Shuster [13] and emphasized the emergence of intermittence with 1 *< µ <* 3 as a generator of 1*/f* noise, namely a noise with the spectrum *S*(*f*) given by

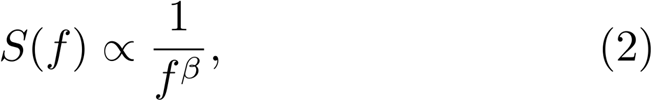

with *β* = 3 − *µ* for 2 *< µ <* 3 and *β* = 0 for *µ >* 3. When the birds fly in different directions making the global velocity of the swarm vanish, a nucleation process starts again making the whole swarm fly in a new direction with no correlation with the earlier flying regime: the changes of flying direction are renewal events. If *µ >* 3 these renewal events generate the ordinary condition of white noise, due to Eq. (2). If *µ <* 3, Eq. (2) signals a departure from the condition of ordinary statistical physics, thereby leading us to the definition of crucial events adopted in this paper. We define *crucial events* as the renewal events when the condition 1 *< µ <* 3 applies.

Crucial events are not confined to the swarms of birds and are a general property of criticality-induced cognition. In the case of brain dynamics [14], ElectroEncephaloGram (EEG) recordings are found to be characterized by Rapid Transition Processes (RTPs) sharing the same properties as that of crucial events [15]. The region *µ >* 3, generating white noise, is virtually indistinguishable from the condition of incompressible randomness [16], corresponding to a lack of cognition. We notice that [17] human brain generates crucial events with *µ* = 2. This condition, according to Eq. (2), corresponds to a perfect 1*/f* noise. Moreover, in general healthy neurophysiological processes also generate the condition of ideal 1*/f* noise which is a marker of maximal resilience. Diseases [18] or difficult tasks [19] may force neurophysiological processes to significantly depart from this ideal condition while still remaining in the crucial regime of *µ <* 3.

We believe that crucial events can be realized spontaneously by the complex systems along the recent lines of the work of Ref. [20–22], advocating Self-Organized Temporal Criticality (SOTC). SOTC is a theory explaining how a system of interacting units may spontaneously generate temporal complexity. It is a form of SOC focusing on crucial events. Currently, the theoretical approach of SOTC is limited to *µ <* 2, but we make the plausible conjecture that this form of spontaneous organization may be extended to the whole crucial regime of *µ <* 3.

In this paper we study an experimental recording of biophoton emission to show that the first phase of the germination of seeds yields crucial events. Nearly a hundred years ago the Russian biologist A. Gurwitsch [23] found that a weak ultra-violet (UV) radiation comes out from the living tissues and influences the mitotic activity of the neighboring tissues. However, despite the confirmation of Gobor [24], this interesting result has been forgotten by the scientific community for many years. With the improvement of the methods to detect weak level of radiation and the theoretical progress in quantum optics, there has been a renewed interest in this phenomenon with the works of Colli and Facchini [25, 26] in the 50s and F.A. Popp [27] in the 80s. These results generated the interpretation of cognition as possible manifestation of quantum coherence [28, 29]. Although the goal of this paper is the detection of crucial events in the process of germination of seeds, it opens up the road to a new way of revealing the possible contribution of quantum coherence to the cognition of living systems, the germinating seeds in this case.

Biophotons are an endogenous production of ultraweak photon emission in and from cells and organisms, and this emission is characteristic of living organisms. This emission is completely different from the normal bioluminescence observed in some simple as well as complex organisms. For example, bioluminescence is at least 1000 times more intense than the biophotons emission. The main characteristics of biophotons measured up to now are the following: the total intensity of the emission goes from several to hundred photons/sec per cm^2^ surface of the living system with a spectral intensity that seems to be quite flat within the energy range between 200 and 800 nm. After any type of stress (chemical agents, excitation by white and/or monochromatic light, temperature), the emission increases by almost a factor of ten and relaxes to the normal values quite slowly following a power law. Finally the photocount statistics that account for the probability of having N photons within some time interval seems to follow a Poissonian distribution. Despite the wealth of experimental phenomenology, the questions of what biophotons are, how they are generated and how they are involved in life are still open.

As pointed out earlier, biophotons play a fundamental role for the cell to cell communication [30, 31]. Recent research papers are using the cell to cell communication through biophotons to shed light into the brain which is usually thought to be the most complex and most intelligent network [32–34]. We make the assumption that a germinating seed is an intelligent complex system and that its communication with the environment, including other seeds, rests on the emission of biophotons. To experimentally demonstrate its intelligence, we analyze the statistics of the bio-photon emission and we show that in the first phase of the germination process *µ <* 3. We find, however, that complexity is a mixture of the dynamics of crucial events and infinite memory that we interpret as a form of quantum coherence. In the late phase of germination, quantum coherence becomes predominant.

The outline of this paper is as follows. In Section II, we describe the experiment conducted to reveal the biophoton emission. In Section III, we illustrate the method of statistical analysis. In Section IV we illustrate the properties of crucial events revealed by the statistical analysis of the photon emission. We devote Section V to discussing the possible role of quantum coherence in the late phase of the germination process. This may become an important goal for future investigation.

## II. DESCRIPTION OF THE EXPERIMENT

Our experimental set up is comprised of a germination chamber, a photon counting system and a turnable filters wheel. See Fig. 1 for details. The photon counting device is a Hamamatsu H12386-110 high-speed counting head that can be powered at low voltage, just +5 V of power supply. The phototube is sensible in the wavelength range between 230 to 700 nm with a small peak at 400 nm. An ARDUINO board driven by a PC with Lab-View program is used for data acquisition and to control the experiment. The acquisition time window is fixed at 1 second and with this time the whole system has a dark current of about 2 photon/sec at room temperature. Seeds are kept in a humid cotton bed put on a petri dish. In this experiment, the sample is formed by 75 lentil seeds. This number has been chosen to have a good signal to noise ratio. A turnable wheel holding a few long pass glass color filters is placed between the germinating seeds and the detector. The wheel has eight positions. Six are used for the color filters, one is empty and the last one is closed with a black cap.

**FIG. 1:**
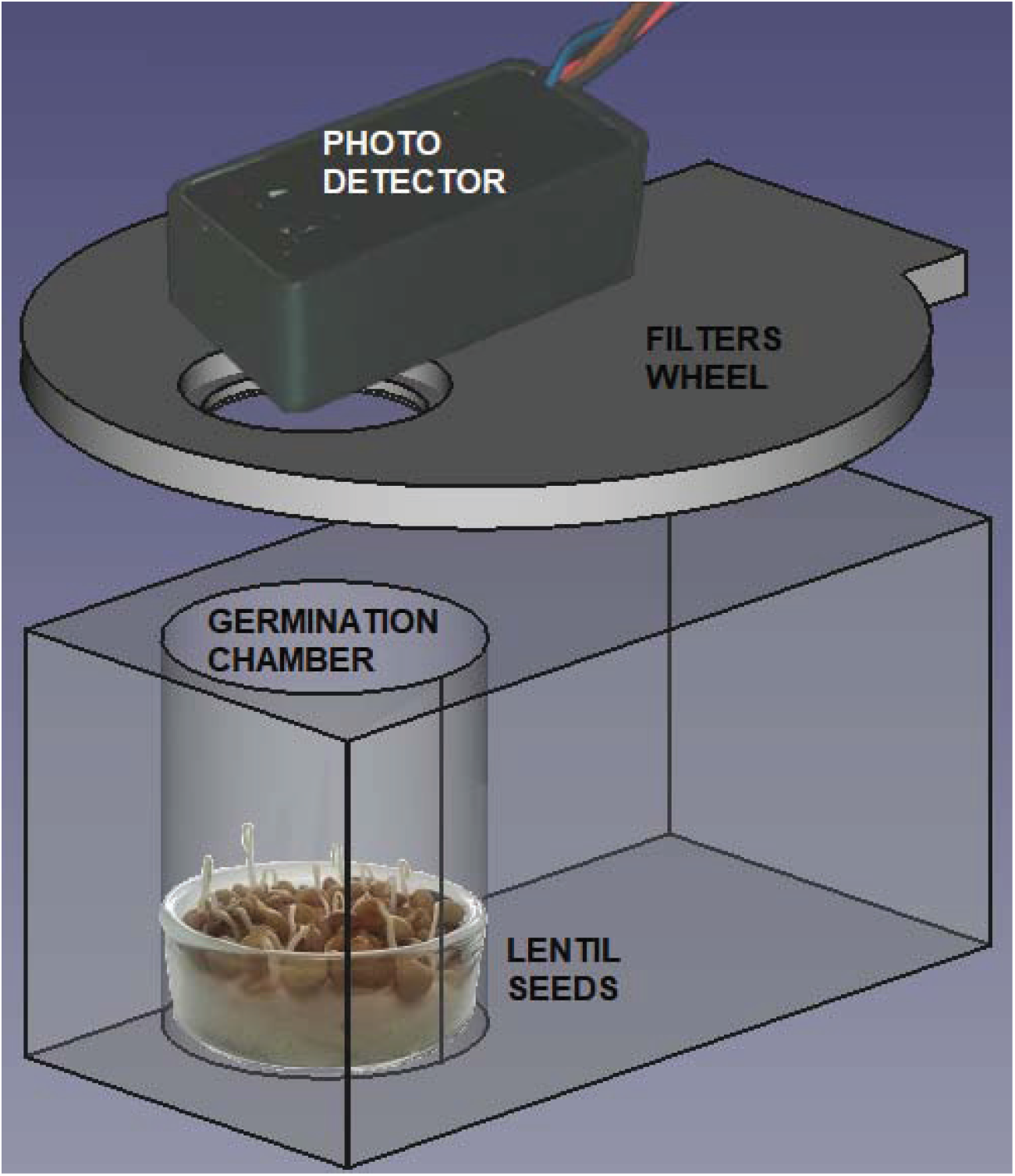
Technical design of the experimental setup used in our experiment. The photon-counting system consists of a Hamamatsu H12386-110 counting head. The germination chamber is built with black PVC to avoid any contamination of the light from outside.

Here, we only use and report the data coming from the empty and black cap positions. This way, we have the total emissions within the visible energy range and the dark counting of the phototube. The wheel stays in each position for one minute. This means that it returns in the same position after 420 seconds plus the time needed to change from one position to the other, a total of 443 seconds. This is the reason why the data here have a bunch-type structure. After one minute of measurement, there are 443 seconds of no data. Without any seeds or germination, there is a monotonic decrease of photon emission which arrives in few hours to the value of the electronic noise. This emission tail comes from the residual luminescence of the materials which is a consequence of the light exposure of the equipment while setting up the experiment. The comparison between the counting in the dark condition (red points) and the signal observed with the germinating seeds (green points) are reported in Fig. 2. The data are related to an acquisition time of almost three days after the closing of the chamber. To clarify the behaviors of the different data sets, we also report the number of photons per second coming from the raw data averaged over one minute.

**FIG. 2:**
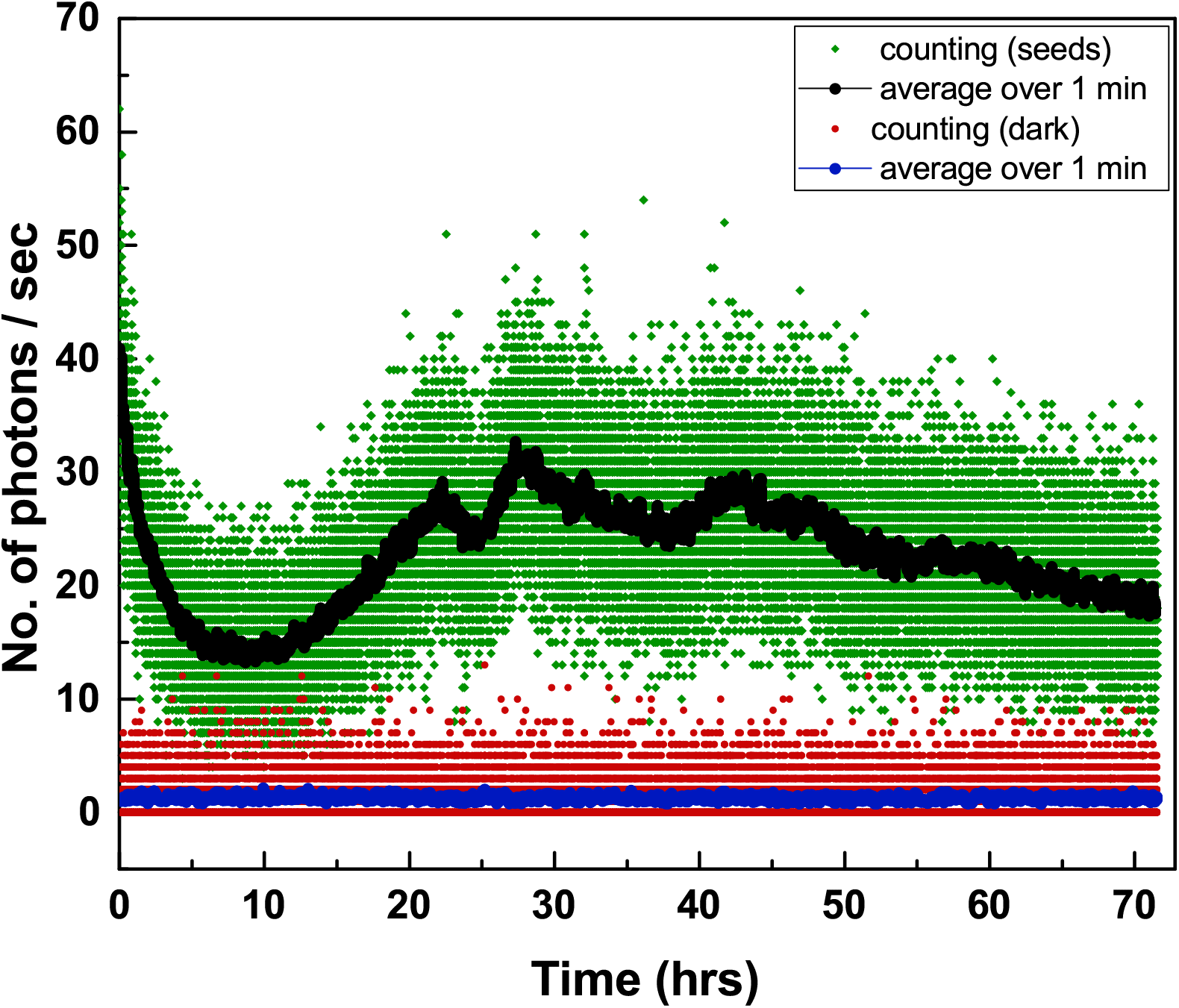
Here we show the comparison between the signal generated by the germinating seeds and the signal in the dark condition. The raw data are the green (seeds) and red (dark count) points. The black and blue curves are the raw data averaged over one minute. In other words, these curves are the countings per second which are averaged over one minute.

Residual luminescence of the whole equipment goes down in few hours up to the time when the lentils start germinating and the ultra-weak (UW) biophoton emission is strong enough to be detected. It is interesting to note how the UW biophotons emission changes during the time showing strong oscillations perhaps due to the different conditions of the seeds during the germination process. The initial behavior is dominated by the luminescence generated by the humid cotton bed. The germination-triggered UW bipphoton emission emerges after about ten hours of closing the chamber and then becomes dominant leading to a saturation effect corresponding to a virtually stationary emission. The signal is well above the electronic noise for the whole time period.

This evolution of the signal is related to the spontaneous evolution to criticality described by Fig. 3 of Ref. [20] where the initial mean field has the value of −1. The mean field then, as an effect of self-organization, moves through a fluctuating transient towards a condition of weaker fluctuations close to the maximal value of 1 signaling the achievement of criticality. Fig. 3 of this paper suggests that water may activate a process of the same type, although criticality associated to germination in the long-time regime may be replaced by a different process characterized by the birth of roots.

**FIG. 3:**
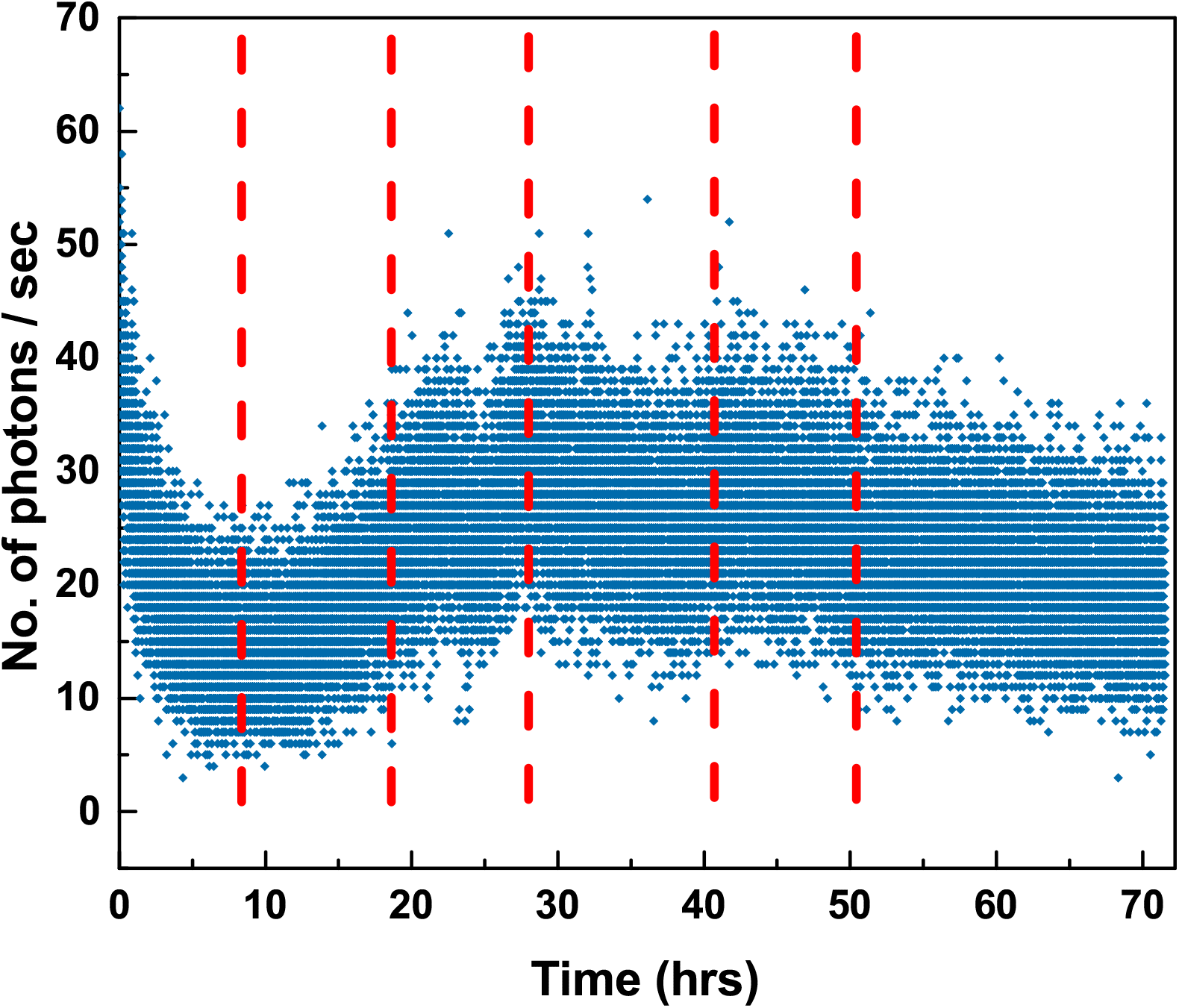
Number of photons emitted during the germination of lentils. The red dashed lines represent different regions during the germination process.

## III. STATISTICAL ANALYSIS

### A. Introduction to the method of diffusion entropy

The method of Diffusion Entropy Analysis (DEA) was introduced in 2001 [35]. It is based on converting the experimental time series under study into a diffusional trajectory, as follows

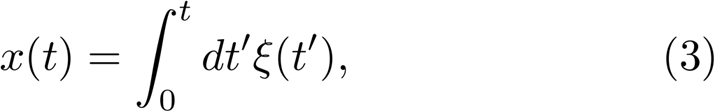

where *ξ*(*t*′) is the time series under study. In this case, *ξ*(*t*′) = *n*(*t*′), where *n*(*t*′) is the number of photons emitted at time *t*′. To make the statistical analysis, we convert this diffusional trajectory into many realizations so as to make it possible to do an ensemble average. These realizations are denoted by

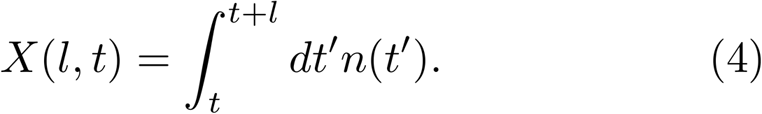

Note that *l* cannot go to infinity because we worked with time series of length *M*. The largest value of *l* is *M − t*. These diffusion trajectories describe the departure from the origin, *X* = 0, of random walkers sharing the property of being at the origin at time *l* = 0. Making *M* very large we created a number of realizations so big as to be able to evaluate the probability distribution density *p*(*X, l*) with an accuracy large enough as to fit the assumed scaling equality

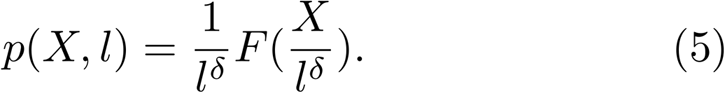

This is a fundamental equation of diffusion processes. Its meaning is as follows: we observed the process at time *l* and at a larger time *L*. The distribution density *p*(*X, L*) was expected to be broader than the distribution density *p*(*X, l*). Of course, since the number of walkers is conserved, the intensity of *p*(*X, L*) is smaller than the intensity of *p*(*X, l*). Let us squeeze the space *X* by changing the values *X* into *X^I^* = *rX*, where *r* = (*l/L*)^*δ*^) and let us amplify the ordinate scale by multiplying *p*(*X, l*) by 1*/r*. When the scaling condition is realized, the distribution density at time *L*, after the abscissa squeezing and amplification of the ordinate, turns out to be identical to the distribution density at the earlier time *l*.

To find *δ* we evaluate the Shannon entropy of the distribution density *p*(*X, l*) given by:

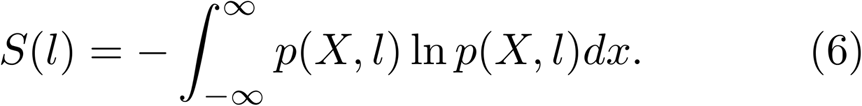

We plug Eq. (5) into Eq. (6). With an easy algebraic calculation we get

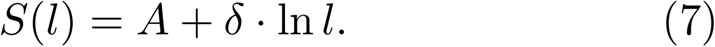

where A is a constant. Thus, DEA allows us to measure the scaling *δ*. In the linear-log representation of *S*(*l*), the scaling *δ* is the slope of the straight line. In the case of complexity, the scaling *δ* is expected to depart from the condition of ordinary diffusion where *δ* = 0.5.

### B. Diffusion Entropy with stripes

How is *δ* ≠ 0.5 related to crucial events? This important issue was discussed in Ref. [36] which devotes a special attention on how to turn the crucial events into a diffusion process. In this paper, we use the DEA of Ref. [37] which adopts the method of stripes to fully benefit from the connection between crucial events and diffusion established by Ref. [36]. In this section, we explain the method of stripes.

First of all, we have to establish a connection between the scaling *δ* and cognition which is interpreted as computational compressibility. The time distance between two crucial events, given by the distribution density of Eq. (1) with the inverse power law index *µ*, is related to the parameter

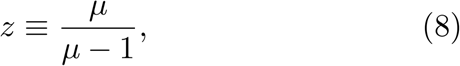

which can be used to define the Kolmogorov-Sinai entropy, *h*_*KS*_, given by [38]

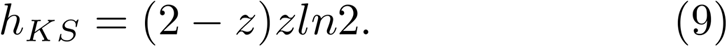

The laminar regions between two consecutive crucial events are filled with symbols, either 1 or 0, randomly selected. The sequence of these symbols generates a time series with a compressibility defined by Eq. (9). The value *z* = 1, corresponding to *µ* = ∞, is the condition of total incompressibility, making *h*_*KS*_ = *ln*2. As discussed in [16], moving from *µ* = ∞ to finite values of *µ* has the effect of making the sequence compressibile, namely, it makes it possible for us to reproduce the time series with a finite number of computer instructions smaller than the size of the time series. The compressibility becomes stronger when *µ <* 3. Notice that at *µ* = 2, *z* = 2, *h*_*KS*_ vanishes. Korabel and Barkai [39] generalized the definition of *h*_*KS*_ so as to explore also the region *z >* 2 (*µ <* 2). The computational cost in the case *µ >* 2 increases linearly in time while in the case *µ <* 2 it increases as *t^α^*, with *α* = *µ* − 1. As a consequence, using *t^α^* rather than *t* for the definition of *h*_*KS*_, Korabel and Barkai [39] showed that *h*_*KS*_ for *z >* 2 becomes an increasing function of *z*. Thus, the generalized *h*_*KS*_ has its minimum value at *µ* = 2 signaling the condition of maximal intelligence shared by the human brain [17]. The Kolmogorov compressibility which implies the existence of cognition is converted into anomalous diffusion using three walking rules. The first rule is that the random walker makes a step ahead by a fixed quantity when a crucial event occurs. The second rule, called velocity rule, is that the random walker moves with constant velocity in the positive or negative direction according to the tossing of a fair coin. In the third rule, called Continuous Time Random Walk (CTRW) rule [36], the random walker jumps only when a crucial event occurs making a step of fixed intensity either ahead or backwards with equal probability. According to Ref. [36], studying the connection between scaling and walking rule, the first rule yields

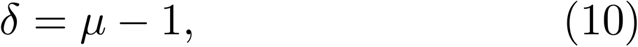

for 1 *< µ <* 2,

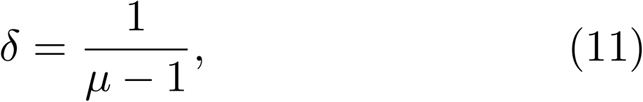

for 2 *< µ <* 3, and

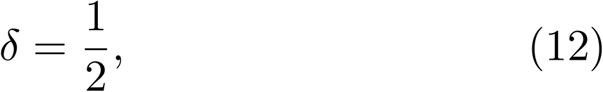

for *µ >* 3. The first rule was shown to afford the fastest scaling in the whole range *µ >* 1 [36] leaving open its competition with the second rule for *µ <* 2, where DEA is affected by big inaccuracies if the second rule is adopted.

After establishing the connection between scaling and walking rule, it is easy to explain to the readers the convenience of adopting DEA with stripes, as done in Ref. [37]. We use the experimental time series *ξ*(*t*), *n*(*t*) in the case of this paper, to define events either crucial or not crucial as follows. We divide the ordinate axis *ξ* into bins of size *s* and record the times at which *ξ*(*t*) makes a transition from one bin to one of the two nearest neighboring bins, including the vertical transitions through many bins in the case when we use bins of small size. This makes it possible for us to create a walking trajectory *x*(*t*) based on the first rule, the walker always making a jump ahead. Then, we use DEA to define the scaling *δ*.

The adoption of DEA with stripe is expected to yield a scaling *δ* that is either equal or smaller than the scaling *δ* obtained with no stripes. This is so because the analysis without stripes generates anomalous scaling *δ >* 0.5 as a result of either Stationary Fractional Brownian Motion (SFBM) or Aging Fractional Brownian Motion (AFBM). It may also generate anomalous scaling generated by a combination of SFBM and AFBM. The scaling of SFBM may be larger than the scaling of AFBM and the adoption of stripes filters the anomalous contribution generated by non crucial events including the non crucial SFBM events. Note that AFBM is the form of memory generated by crucial events,

It is important to stress that the method of stripes can be adopted to establish if cognition is due to criticality or due to quantum coherence [40]. This difficult issue will be discussed in Section V. Here, we limit ourselves to point out that if cognition is due to criticality and *µ >* 2 then the connection between scaling and crucial events is given by

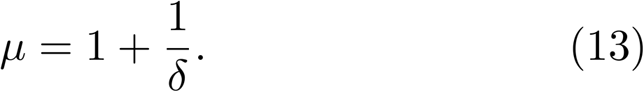

## IV. RESULTS

The results of this Section refer to three questions: (a) Does *µ* change with time throughout the germination process? (b) If not, what is the value of *µ* that represents the intelligence of the germination process? (c) Does the analysis reveal a significant statistical difference between the dark and the germinating state so as to make it plausible to conjecture that cognition emerges during the germination process?

Fig. 3 shows the photon emission from the start of the experiment until the germination process ends with the six regions (separated by the red dashed lines) referring to different stages of analysis. Applying DEA in these different regions, we found the scaling *δ* which is given by Eq. 7 and the corresponding *µ* according to Eq. 12 reported in the Table I. The errors given by the mean squared error (MSE) in Table I are numerical corresponding to the line of best fit when determining *δ*.

**TABLE I:** The scaling µ using DEA in the six different regions of the time series.

| DEA Scaling |  |  |  |  |  |  |
| --- | --- | --- | --- | --- | --- | --- |
| without stripes |  |  |  | with stripes |  |  |
| Region | $\delta_1$ | MSE | $\mu_1$ | $\delta_2$ | MSE | $\mu_2$ |
| 1 | 0.773 | 5.8E-4 | 2.293 | 0.596 | 5.3E-4 | 2.676 |
| 2 | 0.796 | 2.7E-4 | 2.254 | 0.558 | 8.5E-4 | 2.789 |
| 3 | 0.736 | 3.9E-4 | 2.358 | 0.526 | 1.5E-3 | 2.897 |
| 4 | 0.737 | 1.1E-3 | 2.355 | 0.496 | 1.2E-3 | 3.016 |
| 5 | 0.694 | 7.9E-4 | 2.440 | 0.509 | 5.07E-4 | 2.960 |
| 6 | 0.725 | 7.5E-4 | 2.377 | 0.504 | 1.0E-3 | 2.982 |

We see from Table I that the analysis with no stripes yields a scaling significantly larger than the scaling obtained from DEA supplemented by the use of stripes. This indicates that although crucial events exist in the first three temporal regions, the real value of *µ* detected by stripes is much closer to *µ* = 3, the border with the region of ordinary statistical physics, with no complexity. For the last three temporal regions we cannot rule out the possibility that no crucial events exist and that the anomalous scaling, with a significant departure from the ordinary scaling *δ* = 0.5, is due to SFBM. The authors of Ref. [40] demonstrated that the dynamical origin of SFBM is a generalized Langevin equation that is derived using the formalism of quantum mechanics [41], thereby suggesting that SFBM may be a manifestation of quantum coherence.

Fig. 4, referring to region 1, shows how the results of Table I are obtained. Using the concept of intermediate asymptotics [42], we evaluate the slope of *S*(*l*) in an intermediate region between the region of short values of *l* and the region of large values of *l*. The region of short values of *l* is a region of transition to complexity. The region of large values of *l* is either a region of transition to ordinary statistics [20–22] or a region where the scarcity of events makes the evaluation of complexity inaccurate. The errors (given by MSE) of Table I measure the inaccuracy associated in determining the intermediate asymptotics.

**FIG. 4:**
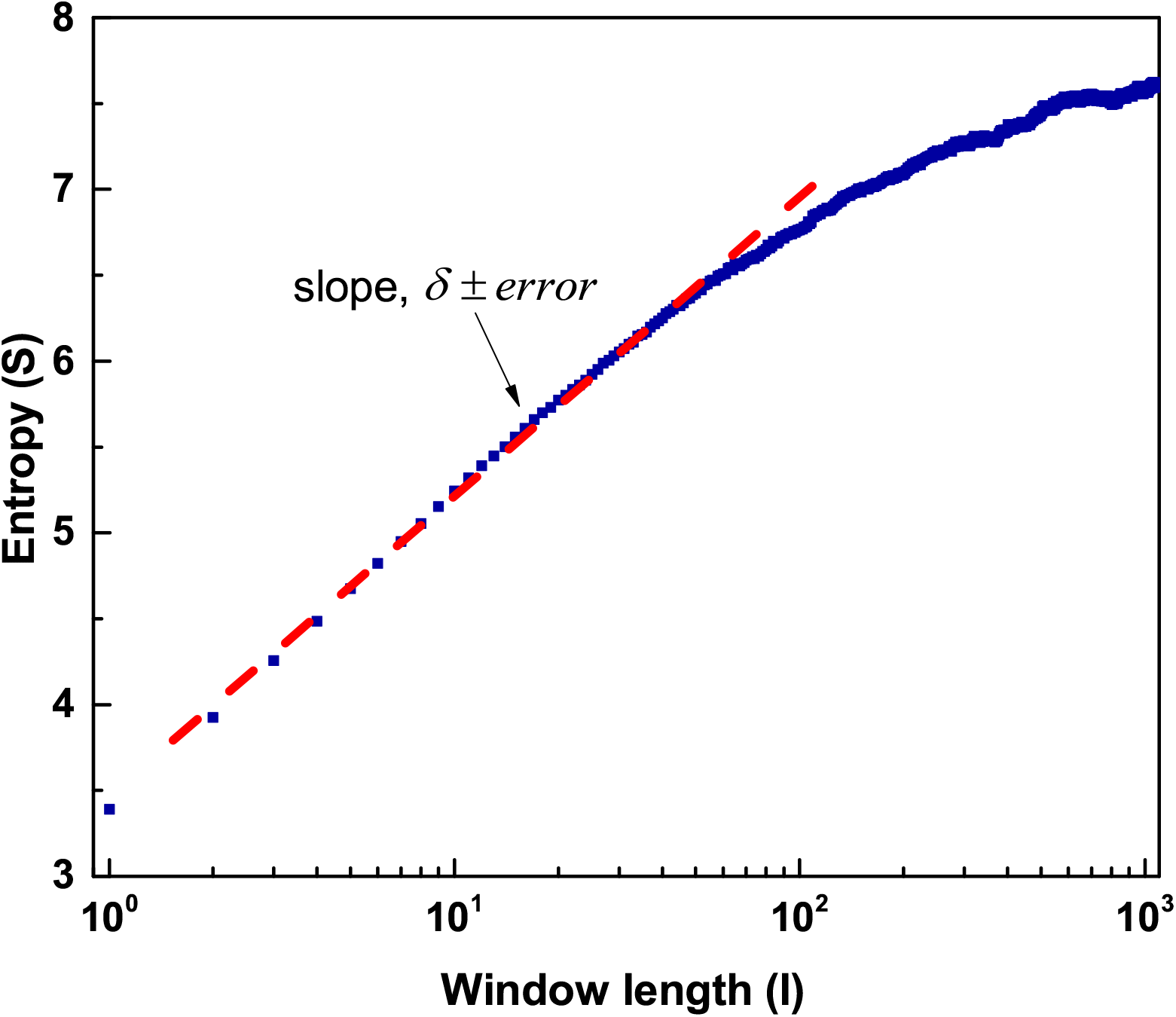
DEA on region 1.

Fig. 5 illustrates the results of our attempts at answering question (a). The results of this figure are obtained by examining portions of different length *L* of the experimental sequence. The starting point of these different portions is always the beginning of the experimental sequence. Increasing the length *L* has the effect of improving the accuracy of the evaluation of *µ*, yielding *µ* ≈ 2.2, however, *µ* is largely independent of the length of the time series *L*. We ignore the region of large values of *l* which may be affected by either the termination of the germination process or by the inaccuracy due to the scarcity of crucial events.

**FIG. 5:**
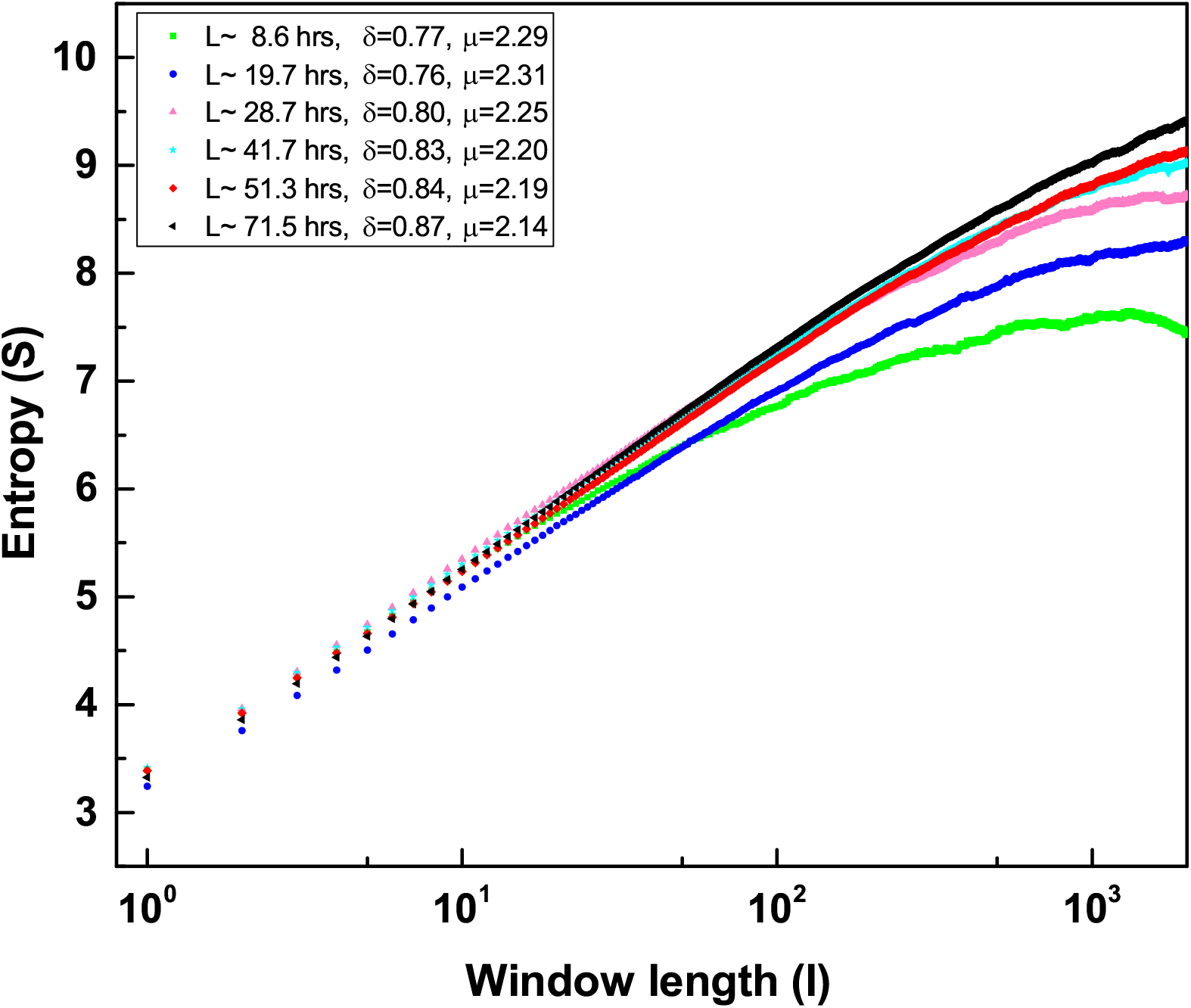
DEA without stripes with changing length L. Different curves illustrate *S*(*l*) for time series of different length, obtained observing experimental data from the time origin up to *L*.

We may answer questions (a) and (b) with the results of DEA supplemented by stripes. The resulting values of *µ* do not significantly change with time and remain in the region of *µ* significantly smaller than 3 which is compatible with the temporal complexity values of neurophysiological processes.

As far as the question (c) is concerned, we address this issue using DEA without stripes and with stripes as illustrated in Fig. 6. The results obtained with stripes lead us to conclude that no temporal complexity exists in the dark state. There is, however, a coherent contribution that is much weaker than the coherent contribution emerging in the long-time region of the germination process. It does not conflict with the main conclusion of this paper that a significant deviation from ordinary scaling either coherent or temporarily complex is due to the germination process. We limit ourselves to notice that the dark state is just the electronic noise. The coherence of the dark state is something inherent to the instrument we are using that was never analyzed, to the best of our knowledge, with the DEA method thereby making it difficult to afford a plausible explanation which is out of scope of this paper.

**FIG. 6:**
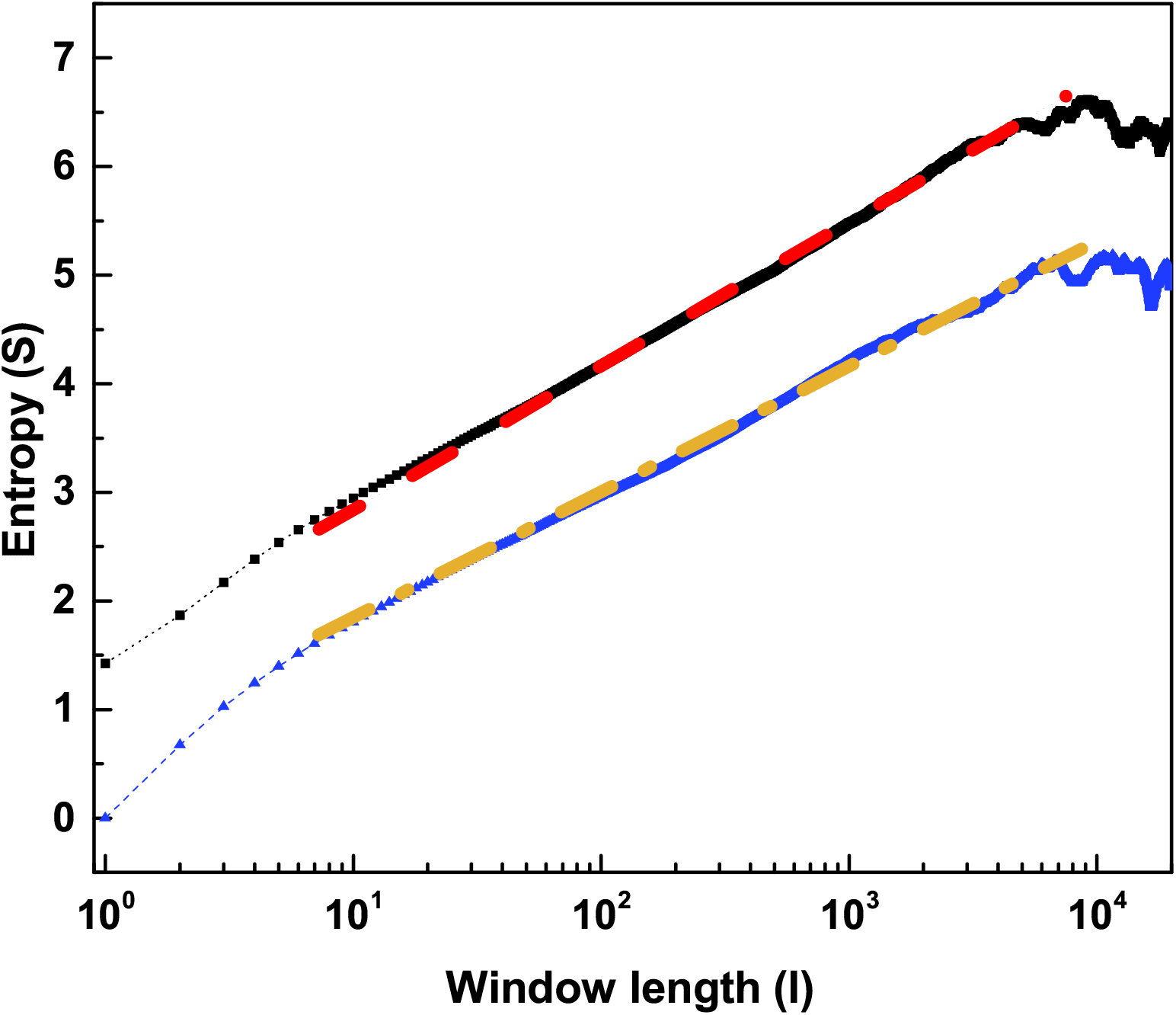
(From top) Black squares represent DEA without stripes applied to the dark state and the corresonding red dashed line shows the fitting with scaling *δ* = 0.575. Blue triangles represent DEA with stripes applied to the dark state and the corresponding orange dashed line shows the fitting with scaling *δ* = 0.508.

## V. DISCUSSION

The method of DEA has been applied in the recent past to a variety of problems such as the application to DNA sequencing [43]. In this case, the DNA sequence is characterized by long laminar regions with purines alternated by long laminar region with pyrimidines. The value of *µ* in those cases is slightly larger than 2. The recent work of Refs. [44, 45] shows that the music of Mozart hosts crucial events with *µ* = 2. It is important to stress that according to the principle of complexity matching [46], two complex systems communicate mainly through their crucial events. This is the reason why the music of Mozart is perceived as being very attractive. In fact, according to Refs. [44, 45] the crucial events hosted by the music of Mozart match the crucial events of the human brain.

Note that the spectrum *S*(*f*) in case when anomalous diffusion is generated by crucial events has the 1*/f* noise form

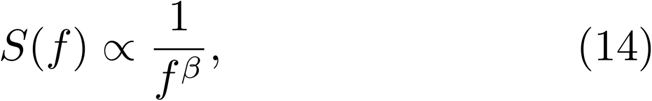

with [47]

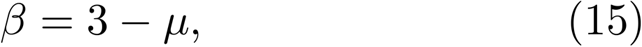

thereby yielding the ideal 1*/f* noise when *µ* = 2.

In the extreme limit where anomalous diffusion is only due to FBM it is known [47] that

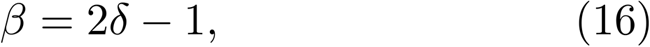

yielding the ideal 1*/f* noise when *δ* = 1.

The results of Table I show that in the first three germination regions a mixture of these two conditions of anomalous diffusion is realized. In the last three germination regions the FBM applies, suggesting the influence of quantum coherence [40]. Therefore, we are forced to adopt a physical interpretation of the transport of information that cannot rest only on the role of crucial events. Nevertheless, we think that this paper affords an important contribution to the revolution occurring in biology [1, 3, 4]. This is so because, first of all, to the best of our knowledge, in the current literature no research work has revealed the existence of crucial events in germination processes. Here, we show that they exist in the first phase of the germination process. Although many authors emphasize the importance of quantum coherence for cognition, including authors believing that biophotons favor cell-to-cell communication, there is no prescription on how to measure and determine the intensity of quantum coherence.

We believe that the results of this paper will lead to future research work where the role of quantum coherence can be established with more precision, by evaluating the spectrum *S*(*f*) and deepening the quantum coherence properties discussed in Ref. [40].

In the literature of biophonons there are indications that the cell to cell communication through light is a plausible communication mechanism, however, there are obstacles to showing this due to the unfavorable signalto-noise ratio of the signal detection by cells [31]. The authors of Ref. [31] discuss the limitation of the famous theory of stochastic resonance. We want to stress that the complexity matching theory of [46] is a much more advanced approach making it possible to establish communication through one single realization rather than an average over an ensemble of realizations, compatible with the joint action of coherence and crucial events. This interpretation may lead to an agreement with that adopted, for instance by Fels [48] where the belief is that the transfer of information and the intelligence itself are based on quantum coherence [28, 29]. In our perspective, however, the origin of cognition is a process of self-organization [20, 21] of the components of the seeds corresponding to water-induced germination. This process of self-organization generates both crucial events and quantum coherence.

These arguments can be used to support our conviction that crucial events exist in the first three temporal regions of the germination process. In the last three regions, the scaling remains significantly anomalous but the main origin of this anomaly is SFBM and according to Ref. [40] this may be a sign of quantum coherence. This leaves room to the supporters of quantum coherence to explain the emergence of cognition. We cannot rule out that the kind of criticality involved by the germination process requires a form of phase transition that is not yet known. It has to be stressed that Mancuso [49] and Mancuso anf Viola [50] use the concept of swarm intelligence with reference to the non-hierarchical root network. Thus, it may be beneficial to supplement their observations noting that the first time region of germination may have to do with the birth of this surprising root intelligence.

## VI. ACKNOWLEDGMENTS

We thank U.S. Army Research Office (ARO) for financial support through Grant W911NF1901. We also thank FQxi for a mini grant. We warmly thank Giuseppe Papalino and Agostino Raco for their help with the building of the experimental setup.

## Notes

#### Summary of Updates

Statistical Analysis section has been revised to include Diffusion Entropy Analysis(DEA) with stripes in order to distinguish between two different sources of anomalous scaling-crucial events with nonstationary correlation function or infinite memory of stationary but non-integrable correlation function. The results section has been updated to include the analysis of the signal with DEA as well as DEA with stripes (updated Table 1). Additionally, Figure 6 has been added to illustrate DEA analysis on the dark count signal in order to compare with germinating seeds.

